# Flicker and Ganzfeld induced visual hallucinations differ in frequency and content

**DOI:** 10.1101/2023.08.08.552408

**Authors:** Oris Shenyan, Matteo Lisi, John A. Greenwood, Jeremy I. Skipper, Tessa M. Dekker

## Abstract

Hallucinatory experiences, defined as perception in the absence of external stimuli, can occur in both pathological and non-pathological states and can be broadly phenomenologically divided into those of a simple and a complex nature. Non-pathological visual hallucinations can be induced experimentally using a variety of stimulation conditions. To assess whether these techniques drive a shared underlying hallucinatory mechanism, despite these differences, we compared two methods: flicker and perceptual deprivation (Ganzfeld). Specifically, we measured the frequency and complexity of the hallucinations produced by these techniques. We utilised button press, retrospective drawing, interviews, and questionnaires to quantify hallucinatory experience in 20 participants. With both experimental techniques, we found that simple hallucinations were more common than complex hallucinations. We also found that on average, flicker was more effective than Ganzfeld at eliciting a higher number of hallucinations, though Ganzfeld hallucinations were longer than flicker hallucinations. There was no interaction between experimental condition and hallucination complexity, suggesting that the increased bottom-up visual input in flicker increased both simple and complex hallucinations similarly. A correlation was observed between the total proportional time spent hallucinating in flicker and Ganzfeld, which was replicated in a retrospective questionnaire measure of experienced intensity, suggesting a shared hallucinatory mechanism between the two methodologies. We attribute these findings to a shared low-level core hallucinatory mechanism, such as excitability of visual cortex, which is amplified in flicker compared to Ganzfeld due to heightened bottom-up input.

## 1 Introduction

Hallucinations are commonly defined as perception in the absence of external stimuli ^1,2^. In the case of visual hallucinations, these false percepts can range from simple - such as geometric forms^3^, to more complex, such as objects, human figures, or landscapes ^4^. As well as occurring in pathological states such as schizophrenia ^5^, Parkinson’s disease ^6^, epilepsy ^7^, migraine ^8^ and vision impairment ^9^, visual hallucinations can also be induced in non-pathological states – through hallucinogens and psychedelics ^10^, and via experimental manipulations such as states of sensory ^11^ and perceptual ^12^ deprivation (known as a ‘Ganzfeld’), and the use of flicker ^13,14^. While experimentally-induced hallucinations are often referred to as ‘pseudo-hallucinations’ due to the participants’ awareness that the hallucinatory experience is not real, the percepts can be very vivid and vision-like^14^. Experimental methods use different means of visual stimulation to induce hallucinations, and it is presently unclear to what extent these different approaches rely on different mechanisms and lead to differences in the nature and complexity of the resulting hallucinations ^14^.

To interrogate this, we compared two experimental methods that differ vastly in the degree of bottom-up stimulation they involve: a salient visual flicker at a frequency appropriate for inducing visual hallucination (flicker), and visual and auditory perceptual deprivation (Ganzfeld). We tested whether increased bottom-up stimulation would alter the frequency and complexity of the hallucinations induced, and also whether the underlying mechanisms of the resulting hallucinations are shared across these methods despite the variations in visual stimulation.

Visual hallucinations can be divided into those of a ‘simple’ or a ‘complex’ nature. Simple hallucinations contain abstract content such as colour alterations, elementary shapes, or geometric patterns. During a systematic study of the subjective effects of mescaline ^15^, Heinrich Klüver outlined four key examples of simple hallucinations including tunnels, spirals, honeycombs and cobwebs. These form constants are cross-cultural and present across a multitude of hallucinatory states such as during psychedelic visual imagery ^16^, migraine ^8^, hypnagogia ^17^ and flicker ^18^, suggesting there may be a shared hallucinatory mechanism of relatively low-level origin underlying their occurrence. Indeed, neural network simulations suggest that form constants could arise from increased excitation in early visual cortices. When aberrant waves of excitation spread across early visual cortices, resulting in stripes on visual cortex, the transformation of these stripes from cortical space to retinal would result in characteristic geometric patterns resembling the aforementioned form constants ^3,19,20^.

Complex hallucinations are those containing figurative elements such as faces, objects, or scenes. The representation of complex hallucinations has been associated with higher-order visual regions ^21–23^. However, in psychedelic visual hallucinations, complex percepts resembling figurative constructs often also incorporate geometric elements, suggesting that there may not be a binary distinction between simple and complex hallucinations, and that lower-level visual areas may be concurrently active with higher-level visual areas during complex imagery ^24^. This could suggest that simple hallucinations arise when activity is strongest in lower-level areas, whereas complex hallucinations arise when this activity travels up the visual hierarchy, either alone or in conjunction with top-down interpretative influences ^13,25,26^. Another explanation is that complex hallucinations are more akin to mental imagery than to veridical vision^27^ – i.e., they are a top-down, or retro-hierarchical process ^26,28,29^.

Our aim was to better understand both the nature of simple and complex hallucinations, and their underlying mechanisms. There are challenges in understanding the nature of these experiences, however, due to the difficulty in both eliciting and measuring hallucinations in a controlled laboratory environment with objective and methodologically sound techniques ^30^. For instance, hallucinations experienced in pathological states can be uncommon and unpredictable - especially in clinical or laboratory settings. This is partly demonstrated by the number of single-subject case studies within the field, suggesting the difficulty of recruiting large numbers of patients experiencing visual hallucinations which can be measured on demand ^31^. Similarly, psychedelics elicit a multitude of changes in conscious experience ^32^ that make it difficult to draw conclusions about their specific effects on the visual system and the hallucinatory state. Greater control over the induction of hallucinations can instead be achieved in a laboratory environment using visual (and auditory) stimulation. Two methods of interest which are known to induce both simple and complex pseudo-hallucinations are high-frequency visual flicker (sometimes referred to as ‘Ganzflicker’), and perceptual deprivation (known as the ‘Ganzfeld’ effect). These methods have different stimulation conditions and have been suggested to rely on different underlying mechanisms^18,33–35^, based on which different predictions can be made surrounding the different types of hallucinations that may be elicited, and their complexity.

The term ‘Ganzfeld’ refers to a state of sensory homogeny or perceptual deprivation (as opposed to sensory deprivation where stimulation is removed, e.g., via blindfold). The technique was first popularised by Metzger in 1930 (ganz = whole; feld = area, field) ^36^, and traditionally refers to the exposure of an individual to homogenous, unstructured, sensory input, which can result in an altered state of consciousness ^12,33^. In recent years, a ‘multimodal Ganzfeld’ has been induced by using halves of ping pong balls securely fastened over open eyes accompanied by light to achieve sensory homogeny ^33,35,37–39^, combined with unstructured auditory stimulation such as white, brown, violet or pink noise to achieve auditory homogenisation ^12,38^. After prolonged exposure to the Ganzfeld, both simple and complex pseudo-hallucinatory percepts have been reported to arise ^12,35^. An fMRI study found reduced connectivity between the thalamus and primary visual cortex (V1) during a Ganzfeld compared to rest, which was attributed to a reduction in bottom-up signalling. When coupled with intact top-down fluctuations in activation, such a reduction in structured bottom-up signalling could cause sensory noise in early visual areas to be mistaken for signal, leading to pseudo-hallucinations ^33^. Such a process may be especially heightened in the Ganzfeld as internally generated percepts do not have to compete with veridical percepts ^26^.

Flicker was first reported to induce hallucinations by Purkinje in 1819. While waving his hand between his eyes and sunlight, he reported ‘beautiful regular figures that are initially difficult to define but slowly become clearer’ ^40,41^. In more recent years, empty-field high-frequency flicker (otherwise known as ‘ganzflicker’ ^14,42^) has consistently been found to produce hallucinations of both a simple ^30,43–45^ and complex nature ^13,14,18,42,46–48^. Flicker-induced simple hallucinations have been proposed to utilise a rhythmic excitatory bias in early visual cortices ^45,49^. In this process, wave patterns of excitation akin to those associated with geometric Klüver-like form constants, are thought to encourage neural entrainment; the synchronisation of the brain’s endogenous neural oscillations (e.g., the alpha rhythm) to an endogenous external stimulus such as the flicker ^18,50 13,34,44^. In line with this, several studies have shown that simple hallucinations are most frequent when flicker is presented at the alpha-wave frequency of approximately 10 Hz ^18,51^ (although hallucinations also occur at other frequencies^13,14^). In the case of flicker-induced complex hallucinations, it has been suggested that basic hallucinatory forms may provide a building block for interpretive top-down influences^13^.

### 1.1 Aims and hypotheses

Given the above differences in both the pattern of visual stimulation and the proposed mechanisms of flicker-and Ganzfeld-induced hallucinations, we sought to directly compare the nature and frequency of the visual pseudo-hallucinations produced by these methods. We hypothesised that the increased bottom-up input from flicker would result in more simple hallucinations than the Ganzfeld (Hypothesis 1). The latter may rely more on visual cortex excitability, which would likely be more variable between individuals. We also predicted that in flicker (Hypothesis 2), there would be more simple hallucinations - such as form constants - than complex hallucinations, due to both their ubiquity across hallucinatory states and their apparent low-level origin. Predictions for simple versus complex hallucinations in the Ganzfeld were more complicated, as outlined under Hypothesis 3 below.

If complex hallucinations are primarily feedforward in origin, i.e. driven by noise travelling up the visual hierarchy (higher than that related to simple hallucinations), then the greater bottom-up signal with flicker should produce more complex hallucinations than the Ganzfeld. The overall ratio of simple to complex hallucinations should nonetheless be similar across the two tasks if the processes underlying the initiation of both types of hallucination are the same, but driven more strongly during flicker (Hypothesis 3a). That is, we would expect more hallucinations overall for flicker than Ganzfeld, with more simple than complex hallucinations for both flicker and Ganzfeld. Alternatively, it may be that complex hallucinations are more driven by top-down processes, similar to mental imagery. If so, then the lack of structure in the bottom-up input in the Ganzfeld may elicit more complex hallucinations – or a higher ratio of complex hallucinations to simple hallucinations - than with flicker, due to a decrease in the hierarchical competition between these inputs and the top-down influences (Hypothesis 3b). This could manifest either with more complex than simple hallucinations for the Ganzfeld (the opposite of our predictions for flicker) or as a proportional increase in complex hallucinations for the Ganzfeld (i.e. where simple hallucinations may be more frequent overall, but proportionally less dominant for the Ganzfeld). Finally, we predicted that if the two means of inducing hallucinations share a common hallucinatory mechanism, then people who are more hallucination-prone and experience more hallucinations from flicker should also experience more hallucinations during the Ganzfeld (Hypothesis 4).

## 2 Materials and methods

### 2.1 Sample

Twenty participants (14 female, six male) with a mean age of 27.5 ± SD 6.66 years (range 18-50 years) completed the study. None of the participants had a history of neurological disorder, including any history of seizure. Participants were recruited through university recruitment systems and word of mouth. All participants provided informed consent. The experimental procedure was approved by the Experimental Psychology Ethics Committee at the Division of Psychology and Language Sciences, University College London. The research was carried out in accordance with the tenets of the Declaration of Helsinki.

The study had a counterbalanced repeated measures design with all participants undergoing 15 minutes of flicker and 25 minutes of Ganzfeld. The duration of the Ganzfeld was determined in line with previous literature ^33,38^, and a pilot study which suggested that it often took participants longer to see pseudo-hallucinatory percepts in the Ganzfeld than flicker. The duration of flicker was chosen based on previous literature ^13,42^ and a pilot study in which some people considered continuous extended flicker above this duration uncomfortable. In both conditions, participants were provided with a keyboard and asked to indicate both the onset and offset of any perceived visual pseudo-hallucination (outlined in *Response measures Button press and drawing)* and to provide a brief prompt of what they saw, which the experimenter noted.

There was approximately a thirty-minute break between conditions to allow for a wash-out period. During this time, participants a) drew their hallucinatory experiences with respect to their given prompts, b) rated their sleepiness and their perception of how much they felt the button press interfered with their experience, c) underwent an open interview, and d) completed two retrospective questionnaires (covered in detail in Supplementary materials - Questionnaires). The study design is given in Figure 1.

**Figure 1:**
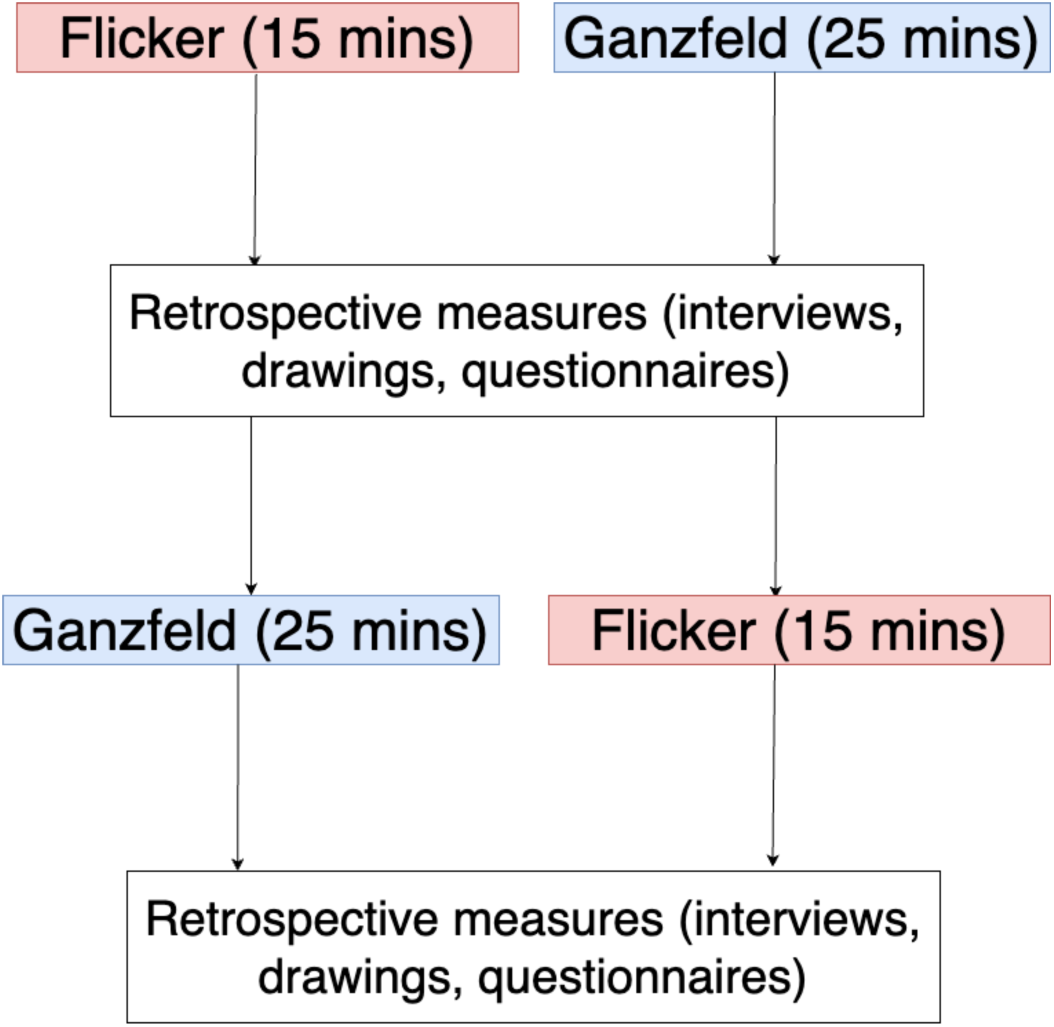
Study design. 20 participants were split into groups of 10. Both groups underwent 15 minutes of flicker and 25 minutes of Ganzfeld. Following each experimental condition, retrospective measures were taken.

### 2.2 Experimental procedures

#### 2.2.1 Instructions

Participant instructions are given in the Supplementary Materials – Detailed participant instructions.

#### 2.2.2 Flicker

Participants were seated in a darkened and soundproofed room, at 70cm from a 32” LED backlit LCD monitor (60hz frame rate; Cambridge Research Systems BOLDscreen). The screen flickered an alternating black and red display at a frequency of 10Hz (as in previous literature^14,42^) for 15 minutes. The frequency was chosen based on multiple studies suggesting that flicker between 8-12hz, and in particular 10hz, is a suitable frequency for eliciting visual pseudo-hallucinations ^18,47,48^. Stimuli were coded using Psychtoolbox-3 running on MATLAB 2021b. Participants first underwent a 30 second practise run during which they saw the flickering stimuli and practised pressing the appropriate buttons and verbalising their experience. If participants were unclear on the instructions, they were allowed another practise run. Auditory brown noise was played through noise cancelling headphones and adjusted per participant to a volume which was comfortable, but loud enough to block out any external sound ^37^.

#### 2.2.3 Ganzfeld

Orange coloured ping-pong balls were halved and taped securely over the eyes using medical tape. The visual field was illuminated by a warm white light (Lumary 24W Smart LED Flood Light). Participants were seated approximately 30cm from the light. Auditory brown noise was played through noise cancelling headphones using the protocol outlined above. While participants were seated, but without their eyes covered, participants practised using the keyboard to report possible hallucinatory experiences.

### 2.3 Response measures

#### 2.3.1 Button press and drawing

For both flicker and Ganzfeld conditions, participants were asked to press the left arrow button on a keyboard when they felt a hallucinatory experience appearing, and to indicate via the right arrow button when the experience had faded away. These button presses provided an indication of the number and the duration of hallucinations experienced by participants. Once the right arrow button was pressed (indicating the percept had faded), participants were instructed to verbalise in as few words as possible what they saw. This was noted by the experimenter. Participants were told that these notes would be used as a prompt for their drawings later. From the button-press data, we counted the number of hallucinations, which were transformed into a rate of hallucinations per minute (i.e. the number of hallucinations divided by 15 for the flicker condition and 25 for the Ganzfeld condition, given their respective durations). We also acquired the average duration of hallucinations, and the total proportional time spent hallucinating (the total summed duration of hallucinations, divided by the length of the experiment).

##### 2.3.1.1 Classification of hallucinations

After the experiment, participants were given their prompts to draw impressions of their hallucinatory experiences – either using paper and coloured pens in earlier iterations of the experiment, or via an iPad (9^th^ generation) in later experiments. These drawings, combined with the associated prompts, gave the experimenter an idea of the nature of the hallucinations experienced and informed their classification as ‘simple’ or ‘complex’.

Simple hallucinations were defined as any descriptions and corresponding drawings of colours shapes, or patterns including characteristic Klüver constants (for instance ‘blob’, ‘blue’, ‘tunnel’, ‘grid’). Complex hallucinations were defined as those with corresponding semantic value (for instance ‘dog’, ‘flower’, ‘galaxy’, ‘face’). When participants felt unable to draw their given prompts (i.e. ‘unsure’, ‘moving’, ‘pulsating’, ‘don’t know’), they were classified as simple hallucinations. We were conservative in our classification of complex hallucinations in particular - concordance between the prompt and the drawing was required to be appropriately classified as a complex hallucination. Twenty such examples are given in Supplementary Table 1. To further compare phenomenology across visual stimulation conditions, we also used the words provided for these prompts for a word frequency analyses.

#### 2.3.2 Questionnaires

Two abridged questionnaires were used to retrospectively assess the subjective experience of participants. We used the questionnaire measures as validation for our button press data.

We used the Altered States of Consciousness Rating Scale (ASC-R or 5D-ASC) ^52^, a well-validated 94 item self-report scale for the retrospective assessment of pharmacological and non-pharmacological induced altered states of consciousness. We chose questions primarily from the Elementary Imagery and Complex Imagery dimensions in line with our research question to interrogate the nature of participants’ subjective experience. Further rationale for the items chosen are described in the Supplementary Materials - Questionnaires, and specific items are shown in Supplementary Table 2.

Though the ASC-R measures visual phenomenology, many of the items are non-specific to hallucination research. Therefore we also utilised questions from the newly developed Imagery Experience Questionnaire (IEQ) ^53^. The IEQ was designed to capture the subjective experience of visual psychedelic experiences more accurately and with greater depth, and many of the items within it are relevant for visual experiences induced by non-pharmacological altered states of consciousness, such as flicker and perceptual deprivation. The dimensions from the IEQ are divided into Complexity, Content and Progression. We only used the Complexity and Progression items. In line with our research question, we carried out analyses pertaining to hallucination complexity by separating items from the Complexity dimension of the IEQ into simple (items 1, 2, 3 and 4) and Complex (items 5, 6, 7 and 8). The relevant items used within the study are in Supplementary Table 3.

#### 2.3.3 Rating scales and open interview

Immediately after the experiment, participants were asked to score their a) sleepiness and b) opinion on how much they felt the button press and talking about their experience during the experiment interfered with their visual hallucinations on a 7-point Likert scale from 0 (Not at all) to 6 (very much so).

Participants also underwent an unstructured, open interview, where they were asked ‘Please tell me anything you feel might be relevant to your flicker or Ganzfeld experience. This could include the visual elements of your experience, but also how you felt. If you feel it is relevant, you can also talk about how tired you were during the experience and how much you feel the button press and discussing your hallucinations during the experiment impacted your experience’. We conducted exploratory word clouds analysis using the words collected per visual stimulation condition.

### 2.4 Statistical analyses

Shapiro-Wilk normality tests were used to assess the normality of our dependent variables. Of our dependent variables, only ASC and IEQ scores were normally distributed. To assess whether simple and complex hallucinations were more likely to occur in flicker compared to Ganzfeld in our frequency data (hallucinations per minute), we used negative binomial distribution mixed effects models with condition (flicker vs Ganzfeld) and complexity (simple vs complex) as fixed-effect predictors alongside an interaction term. We allowed intercepts to vary randomly per participant to accommodate individual heterogeneity, and included an offset term to account for the duration of each respective experiment. We modelled the hallucination frequency data with a negative binomial distribution as they were count data (transformed to frequencies) composed of only non-negative integers with a positive skew and many zeros^54^.

To address our hypotheses, we used contrasts to test for differences in complex and simple hallucinations **across** conditions (Hypothesis 1), and between simple and complex hallucinations **within** conditions (Hypothesis 2). We also checked for interaction effects between condition and hallucination complexity in order to test whether differences between simple and complex hallucinations are dependent on condition (i.e. whether complex hallucinations are more likely to occur in one condition over another) (Hypothesis 3). We report the beta parameters corresponding to these effects and their significance. For our count data specifically, we also supplemented our analyses for Hypothesis 3 with a Chi-squared test to examine differences in numbers of hallucinations based on both experimental condition and hallucination complexity.

We used a similar approach to test for differences in average duration of hallucinations across visual stimulation conditions and for different complexities, and to test for differences in questionnaire scores across these factors. For duration data we carried out gamma mixed effects models with a log link because this captures the distribution of this type of data (non-negative, positively skewed), again taking condition and complexity as fixed-effects predictors, and participant number as a random-effects predictor with varying intercepts. As our questionnaire data (ASC and IEQ) were normally distributed, we used a linear mixed effects model, again with the same model structure.

When any other statistical methods were used, these are noted in the Results section. In brief, for comparisons between two groups, t-tests or (Wilcoxon signed-rank tests in the case of non-parametric data) were used. When testing for correlations in hallucination measures across Ganzfeld and flicker (Hypothesis 4) we used Spearman’s rank correlations for non-parametric data, and Pearson’s correlation for parametric data.

All analyses were carried out in R version 4.3.

#### 2.4.1 Exploratory qualitative analyses

##### 2.4.1.1 Word frequency analysis

We used a word frequency analysis to identify the most frequent words used as interview prompts by participants during the hallucinatory conditions. We tokenised all prompts and removed “stop-words” considered to add no meaning, and therefore to be ignored. We formulated stop-words specific to the study ("screen", "orange", "black", “red”, “Ganzfeld”, “hallucination”, “ping pong balls”, “flashing”, “flicker”) and combined these with the stop_words dataset from the R package tidytext. For each remaining word, we counted the frequency, and the percentage of total word mentions in the condition that this corresponded to.

##### 2.4.1.2 Word clouds

Open interview data was used to create word clouds using the R package wordclouds after the removal of the previously mentioned stop words.

## 3 Results

### 3.1 Individual examples

The plots in Figure 2 show examples of the time series and accompanying drawings and prompts for the hallucinations that two participants experienced in the flicker and Ganzfeld condition. These two participants demonstrate the variability and the spectrum of hallucinatory experience that participants could encounter, ranging from very minor perceptual phenomena to multiple complex hallucinations.

**Figure 2:**
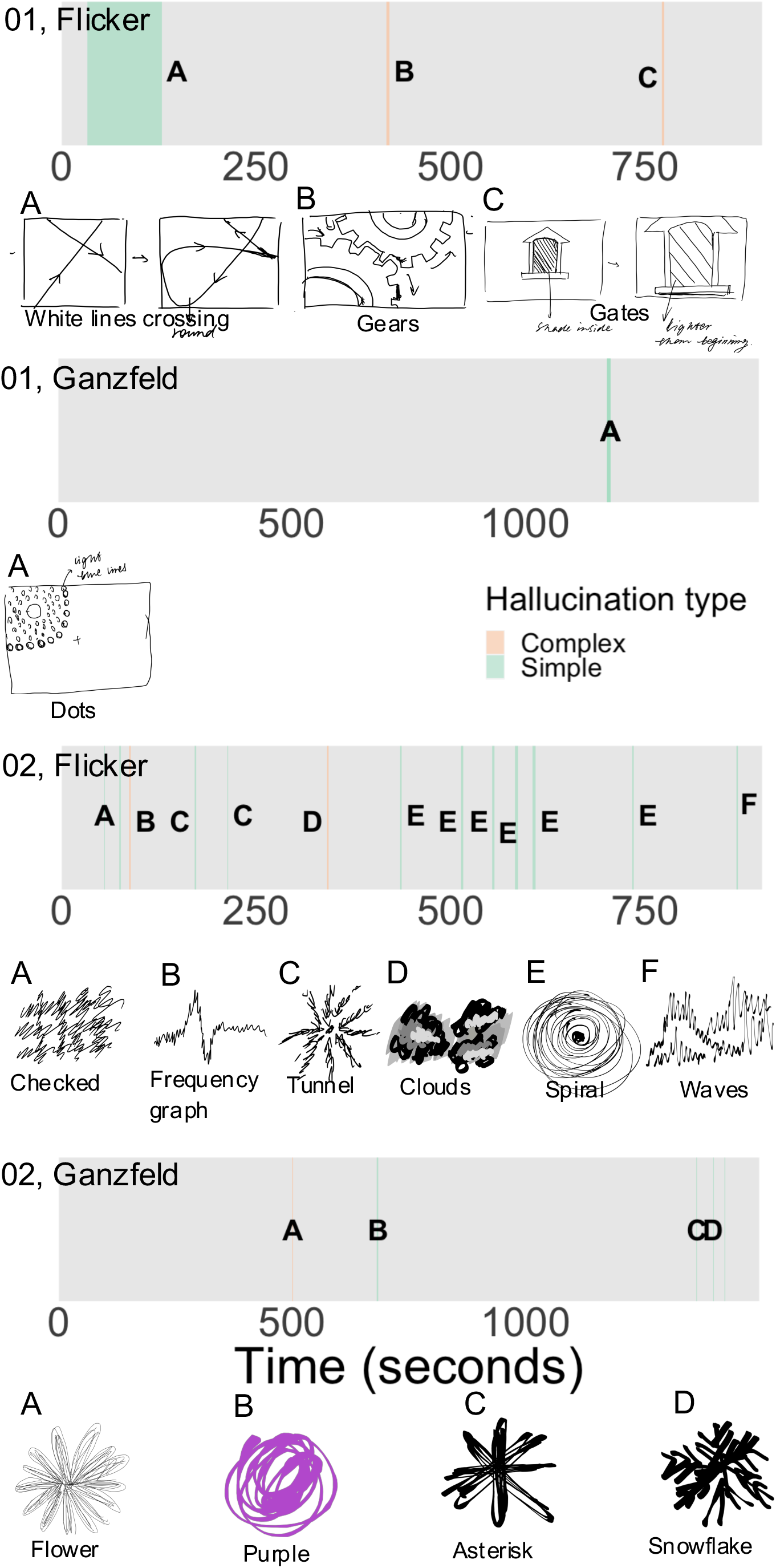
Individual time series for flicker and Ganzfeld for two participants (indicated as 01 and 02). Grey bars depict the time course of each trial, with coloured regions showing the onset and duration of pseudo-hallucinations (green for simple and orange for complex). Hallucinations in each case are marked with letters, with the corresponding drawings that were subsequently produced by participants shown below.

### 3.2 Primary analyses

To provide a generalised overview of the time course of simple and complex hallucinations in flicker and Ganzfeld, we show the probability distributions of simple and complex hallucinations (as defined as the time of a button press onset) in Figure 3. During flicker (A), simple hallucinations tended to have an earlier onset (peaking at 107 seconds) than complex hallucinations (peaking at 519 seconds). This pattern was also present for the Ganzfeld (B), though overall onset times were later than in the flicker condition, with simple hallucinations peaking at 498 seconds compared to 923 seconds for complex hallucinations. Interestingly, simple hallucinations show a bimodal distribution in both the flicker and the Ganzfeld, with a secondary peak at 553 seconds in the flicker and a secondary peak at 1160 seconds in the Ganzfeld.

**Figure 3:**
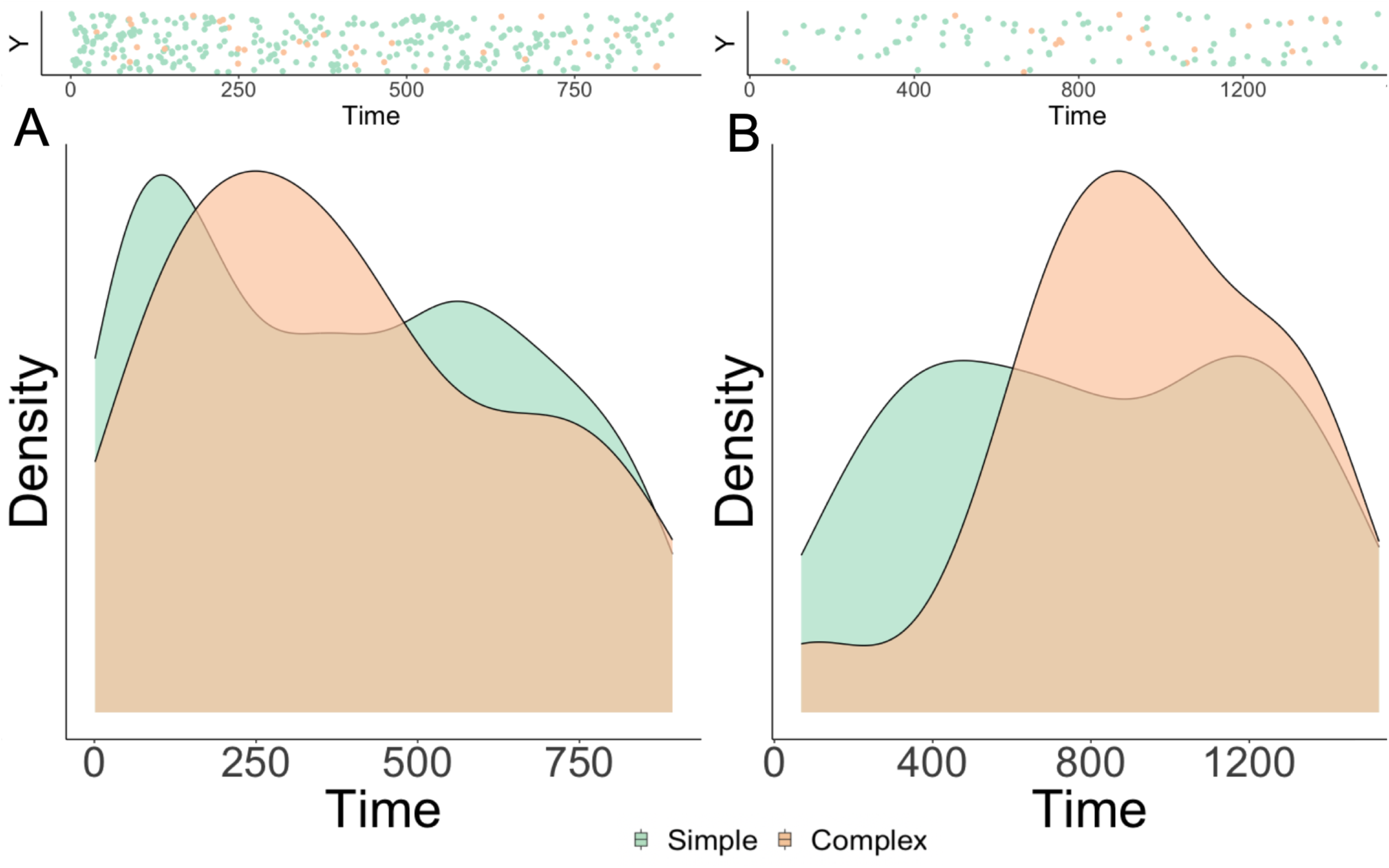
The time course of hallucinations. Scatter plots (shown in upper panels) show the onset of both simple (green) and complex (orange) hallucinations across the time course of flicker (A, 900 seconds) and Ganzfeld (1600 seconds), pooled across participants. Lower panels show the corresponding normalised probability distributions (the probability of a hallucination occurring with each distribution peaking at 1) of simple (green) and complex (orange) hallucinations in flicker (A) and Ganzfeld (B), across the time course of the experimental conditions.

We next compared the frequency of simple and complex hallucinations across flicker and Ganzfeld using negative binomial models (Figure 4A). We show that as expected, within conditions, the rate of simple hallucinations was greater than the rate of complex hallucinations for both flicker (β=2.38, *SE* = 0.36, *Z*=6.57, *p*<0.001) and Ganzfeld (β=1.46, *SE* = 0.40, *Z*=3.67, *p*<0.001). Across conditions, the frequency of hallucinations was higher in flicker compared to Ganzfeld, both for simple (β=1.92, SE = 0.32, Z=5.97, p<0.001) and complex (β=1.01, SE = 0.43, Z=-2.34, *p*=0.019) hallucinations, though there was no significant interaction between condition and complexity (β=0.92, *SE* = 0.53, *Z*=-1.72, *p*=0.09). Descriptively, the ratio of simple to complex hallucinations was 9:1 during the flicker task while it was 5:1 during the Ganzfeld task. We then used a Chi-squared test to assess differences between these ratios, which again showed that there was no significant association between hallucination complexity and experimental condition (*X*^2^ (1, N=436) = 3.14, *p*=0.076).

**Figure 4:**
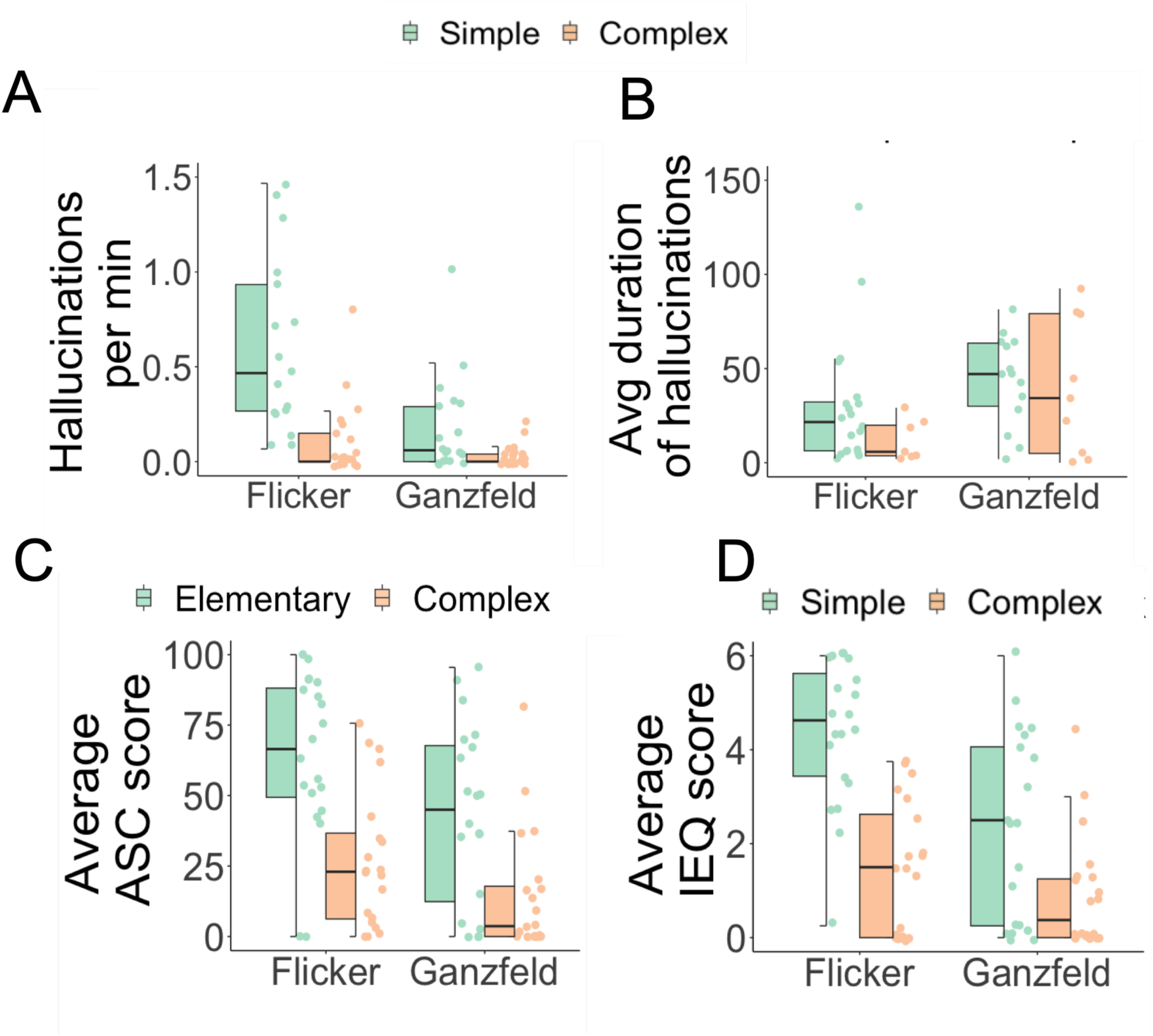
A) The frequency of simple (green) and complex (orange) hallucinations, plotted as the total number of hallucinations divided by the duration of experimental conditions (flicker - 15 minutes; Ganzfeld – 25 minutes) on the y-axis, across experimental conditions (x-axis) B) The average duration of hallucinatory periods in seconds (y-axis) of simple and complex hallucinations across conditions (x-axis) C) Average ASC scores (y-axis) split between Elementary Imagery (green) and Complex Imagery dimensions (orange) across experimental conditions (x-axis) D) Average IEQ scores (y-axis) split between Simple Imagery (green) and Complex Imagery components (orange) across experimental conditions (x-axis). All plots are *N*=20. ASC – Altered States of Consciousness (Rating Scale); Avg – average; IEQ – Imagery Experience Questionnaire; min – minute. The y-axis of plot A has been capped at 1.5 for visualisation purposes; two data points exceeded this value. Similarly, the y-axis of plot B has been capped at 150 for visualisation purposes, two data points exceeded this value.

To summarise, the frequency of experiencing a simple hallucination was greater than that of experiencing a complex hallucination during both flicker and Ganzfeld, and both types of hallucinations occurred more frequently in flicker than in Ganzfeld. Crucially, the likelihood of experiencing a complex hallucination was not significantly higher during Ganzfeld than flicker, although there was a marginal trend towards this effect.

We then tested if the average duration of hallucinations (i.e., the time from start to end button press) varied by content and visual stimulation condition using gamma mixed effects models with a logistic link function (Figure 4B). Across conditions, hallucinations were shorter in flicker compared to Ganzfeld for both simple (β=-1.32, *SE* = 0.11, *T*=-12.53, *p*<0.001) and complex (β=-0.80, *SE* = 0.25, *T*=-3.21, *p*=0.001) hallucinations. Within conditions, there was no significant difference in the duration of simple versus complex hallucinations in the flicker (β=0.14, *SE* = 0.18, *T*=-0.8, *p*=0.42) or the Ganzfeld (β=0.38, *SE*=0.22, *T*=1.7, *p*=0.08) alone. Again, there was no significant interaction between condition and complexity (β=0.51, *SE*=0.27, *T*=-1.9, *p*=0.057). Thus, the average duration of a reported hallucination was longer during the Ganzfeld than the flicker but was not significantly modulated by complexity.

To investigate whether people who reported more hallucinations when looking at flicker also reported more hallucinations during the Ganzfeld, we ran Spearman’s rank correlation analyses. When combining the frequency and duration of hallucinations into a combined measure of overall time spent hallucinating during the visual stimulation session, (total proportional duration time spent hallucinating), we found a positive correlation between experiences during the flicker and Ganzfeld (r_s(18)_= 0.55, p=0.012, Figure 5). The correlation between the frequency of flicker hallucinations and the frequency of Ganzfeld hallucinations was not significant (r_s(18)_= 0.35, p=0.134), and neither was the correlation between the average duration of flicker hallucinations and the average duration of Ganzfeld hallucinations (r_s(18)_= 0.34, p=0.15).

**Figure 5:**
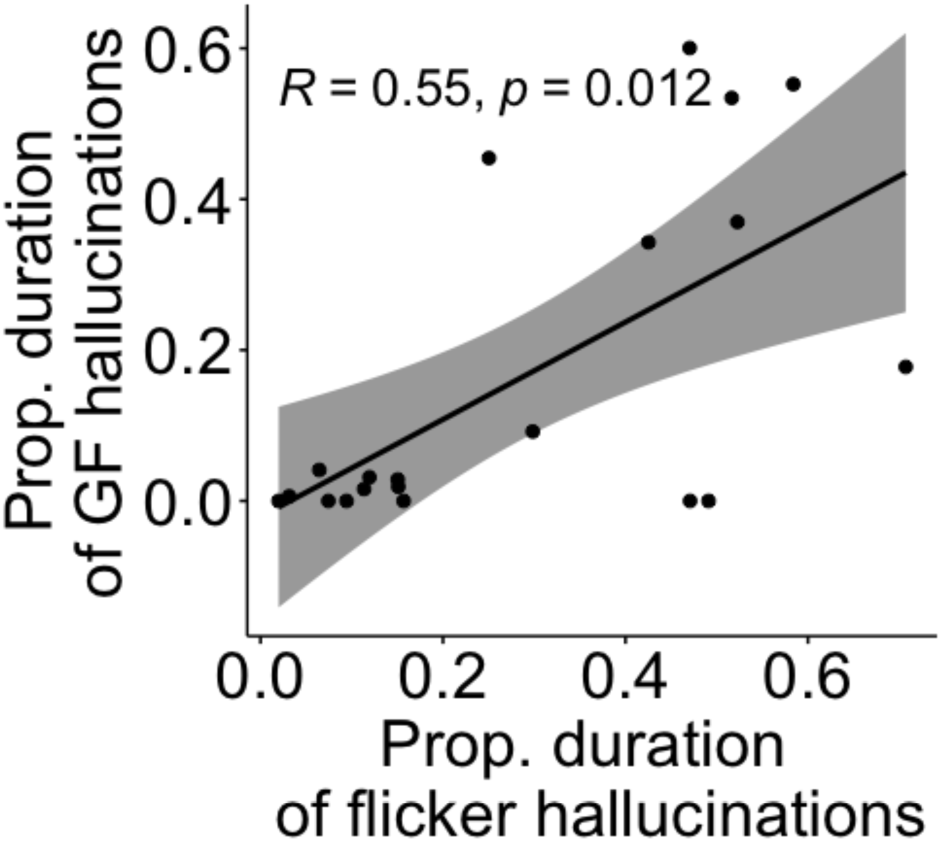
Scatterplot showing the relationship between the total proportional time spent hallucinating (total time spent hallucinating divided by the duration of the experimental condition in flicker (x-axis) compared to Ganzfeld (y-axis) with associated trend line (black) and Spearman’s rank correlation testing (95% CI; grey shading) for *N*=20. GF – Ganzfeld; prop – proportional.

#### 3.2.1 Questionnaire validation

Button presses provide a direct and objective measure of hallucination frequency, onset, and offset. However, they may be affected by non-experiential variables such as the criterion for when to press (i.e., some people may press for a very faint experience, while others may press only for a very vivid experience). Therefore, we validated our button press findings with subjective hallucination intensity as measured through retrospective questionnaires. Using linear mixed effects models, we show across conditions that in the ASC (Figure 4C), Elementary imagery scores were higher in flicker compared to Ganzfeld (β =20.5, *SE* = 7.71, *T*= 2.66, *p*= 0.011). Though Complex Imagery scores were numerically higher in flicker than in Ganzfeld, this was not significant (β = 12.53, SE = 7.71, *T*=1.63, p=0.11). Within conditions, Elementary Imagery scores were greater than Complex Imagery scores in both flicker (β=36.55, *SE* = 7.71, *T*= 4.74, *p*<0.001) and Ganzfeld (β =28.58, *SE* = 7.71, *T*= 3.71, *p*<0.001). There was no significant interaction effect between condition and complexity (β=7.97, *SE*=10.91, *T*=0.73, *p*=0.47). In other words, the ASC scores pertaining to Elementary Imagery were higher in flicker than in Ganzfeld, and ASC Elementary Imagery scores were greater than Complex Imagery in both conditions. Again, the likelihood of higher Complex Imagery scores was not significantly higher during Ganzfeld than in flicker.

We found a similar pattern in the IEQ (Figure 4D). Across conditions, using linear mixed effects models, Simple Imagery scores were significantly higher in the flicker compared to the Ganzfeld (β =2.01, *SE* = 0.41, *T*= 5.01, *p*< 0.001). Again, though Complex Imagery scores were numerically higher in the flicker compared to the Ganzfeld, this was not significant (β=0.59, *SE* =0.41, *T*=1.43, p=0.16). Within conditions, Simple Imagery scores were greater than Complex Imagery scores in both flicker (β =2.89, *SE* = 0.41, *T*= 7.02, *p*<0.001) and Ganzfeld (β =1.41, *SE* = 0.41, *T*= 3.43, *p*=0.011). There was also a significant interaction effect, suggesting that the likelihood of having a higher a Complex Imagery score in the IEQ is higher during Ganzfeld than in flicker. (β=1.48, *SE*=0.58, *T*=2.53, *p*=0.014). We provide a visualisation of a question-by-question analysis for the ASC and IEQ in Supplementary Figure 1.

We also carried out correlations between total ASC and IEQ scores across conditions. We found a positive correlation between average IEQ scores in the flicker and IEQ scores in the Ganzfeld (r_(18)_= 0.35, p=0.036), although the correlation in the ASC was not statistically significant (r_s(18)_= 0.31, p=0.19).

Thus, retrospective questionnaire measures largely replicated the results obtained with button press and hallucination prompts. In addition, we sought to explore whether participants felt that the button press interfered with their experience, e.g. through decreasing the duration of hallucinations - specifically in the Ganzfeld, which may require a higher degree of immersion for hallucinations to occur. ^38^ We did not observe any evidence that the button presses interfered with the occurrence of hallucinations - there were no correlations (p>0.05) between our button-press measures and participants’ perception of the interference of the button press, nor were there any differences between perception of button press interference experience across the conditions (Full details given for analysis in Supplementary materials – Button press interference and sleepiness, V=33, p=1, Supplementary Figure 2B).

#### 3.2.2 Additional analyses

##### 3.2.2.1 Word frequency analysis

A word frequency analysis was undertaken to identify common words within the prompts given by participants during the button-press component of the experiment. The ten most frequently reported words for simple and complex hallucinations in the flicker and Ganzfeld condition are given in Table 1. Examples of the drawings of the most frequent words are given in Figure 6. Interestingly, though there are words of a dynamic nature (i.e. spinning, moving, pulsating) in both conditions, these were more common in the flicker than the Ganzfeld. In addition, there are references to geometric form constants during flicker (i.e. tunnel, Figure 6A, 6B and 6C), whereas the percepts described during the Ganzfeld tend to lack this geometric regularity (i.e. bubbles, blobs, cloud, Figure 6I, 6K and 6L). In addition, despite the warm hued tint of the light stimulation during Ganzfeld, words associated with colour tended to be more cool toned (blue, green, Figure 6I and 6L).

**Figure 6:**
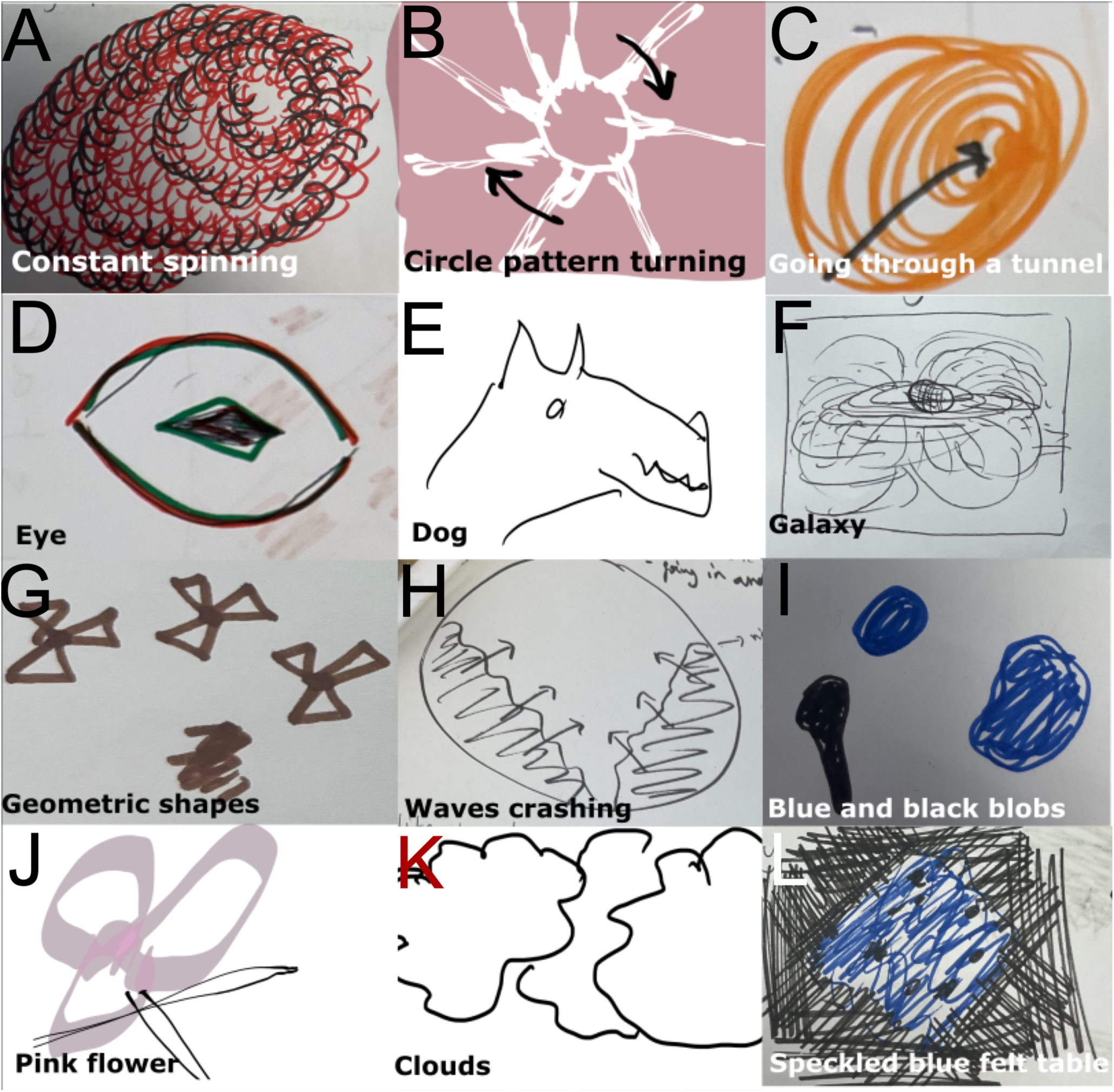
Examples of the most common simple and complex hallucinations as given by prompts given by participants during experimental conditions in flicker (simple – A, B, C; complex – D, E, F) and Ganzfeld (simple – G, H, I; complex – J, K, L). Abbreviated prompts accompany drawings.

**Table 1:** Word frequency analysis illustrating the frequency of words used in the hallucinatory prompts given by participants during flicker and Ganzfeld, split out by hallucination complexity (simple/complex)

| Flicker |  | Ganzfeld |  |
| --- | --- | --- | --- |
| Simple, n(%) | Complex, n(%) | Simple, n(%) | Complex, n(%) |
| Spinning, 26<br>(3.19) | Dog, 6 (6.25) | Shapes, 11<br>(3.34) | Blue, 3 (3.85) |
| Light, 19 (2.33) | Galaxy, 4 (4.17) | Bubbles, 8 (2.43) | Flower, 3 (3.85) |
| Lines, 18 (2.21) | Indistinct, 3<br>(3.12) | Light, 8 (2.43) | Green, 3 (3.85) |
| Moving, 18 (2.21) | Shape, 3 (3.12) | Moving, 8 (2.43) | Cloud, 2 (2.56) |
| Tunnel, 18 (2.21) | Bridge, 2 (3.12) | Pulsating, 7<br>(2.13) | Colours, 2 (2.56) |
| Circles, 17(2.08) | Coming, 2<br>(2.13) | Blue, 6 (1.82) | Hollow, 2 (2.56) |
| Circle, 13 (1.59) | Eye, 2 (2.13) | Lines, 6 (1.82) | Lines, 2 (2.56) |
| Centre, 12(1.47) | Eyes, 2 (2.13) | Waves, 6 (1.82) | Microbe, 2<br>(2.56) |
| Middle, 12 (1.47) | Planet, 2 (2.13) | Blobs, 5 (1.57) | Purple, 2 (2.56) |
| White, 12 (1.47) | - | Dots, 5 (1.52) | Speckled, 2<br>(2.56) |

##### 3.2.2.2 Word clouds

Word clouds created from the open interview data are given in Figure 7 for flicker (A) and Ganzfeld (B). Interestingly, references to form constants are present in the flicker condition (tunnels, patterns, spirals), but not in the Ganzfeld., and there are more references to complex or figurative constructs within the flicker condition, (stars, dogs, hearts, people). Minor perceptual phenomena which could be related to phosphenes (line, round, oval, dots, circle(s)) were present in both conditions. In both conditions, there are frequent references to the dynamic nature of the experience (move(ing), fast, change(d)).

**Figure 7:**
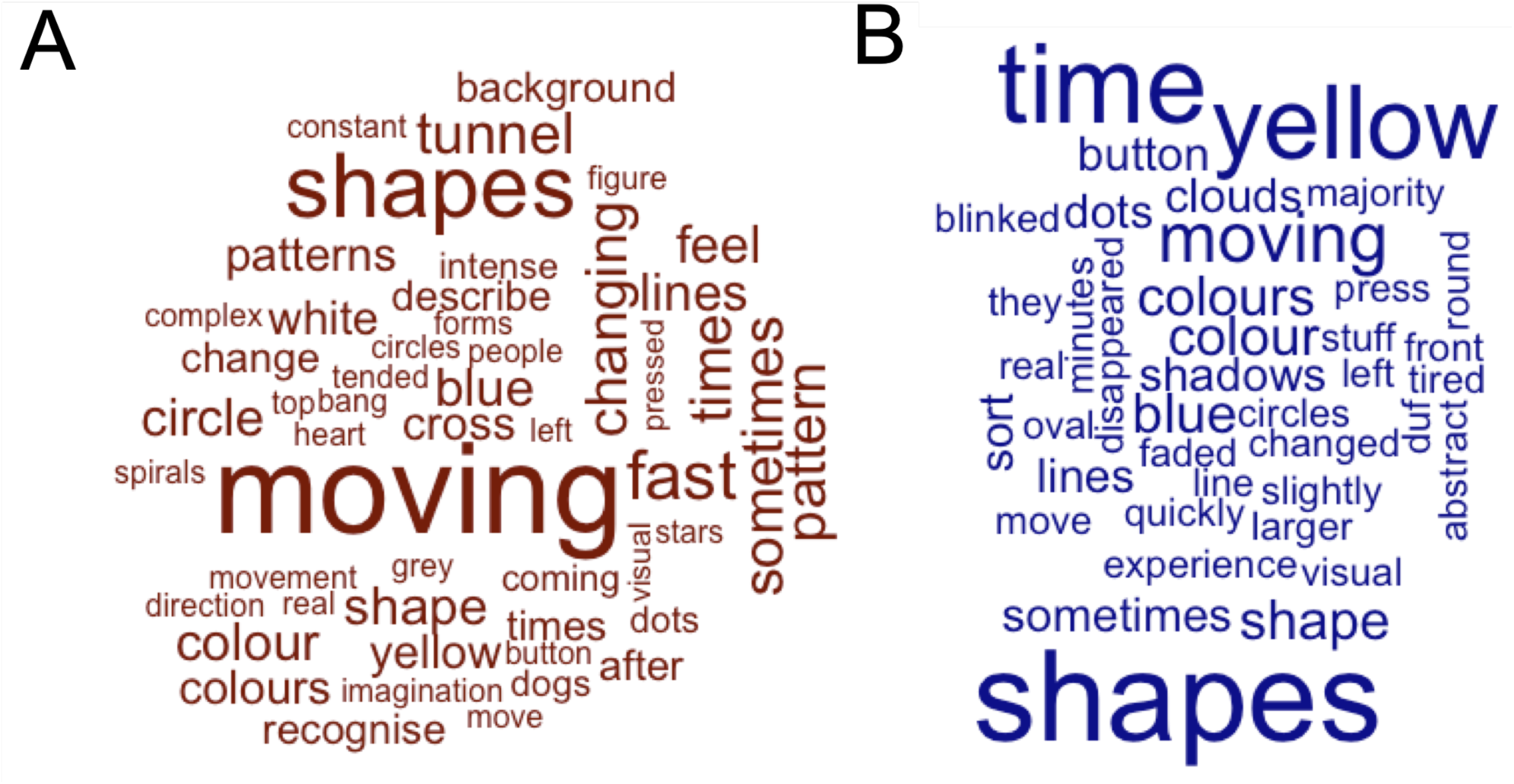
Word clouds for flicker (A, red) and Ganzfeld (B, blue) created from open interview data. Size of words is indicative of frequency of usage in open interview data.

##### 3.2.2.3 Sleepiness and button press measures

We explored relationships between participants’ measures of sleepiness and their perception of how much the button press interfered with their experience. Our rationale for this was that sleepiness may affect proneness to hallucinations – i.e., in hypnagogia^39^, and may vary across the flicker and Ganzfeld condition.

Details for this analysis are given in full in the Supplementary materials – Button press interference and sleepiness. In brief, paired Wilcoxon signed rank tests suggested that participants felt significantly sleepier in Ganzfeld than in flicker (V=0, p<0.001, Supplementary Figure 2A). There was a negative correlation between sleepiness and the total proportional time spent hallucinating (i.e. as people felt more sleepy, they spent less time in total hallucinating) (r_s(36)_=-0.35, p=0.031). No other correlations between button press measures and sleepiness measures were significant (p>0.05), so this factor is unlikely to have driven all task differences.

##### 3.2.2.4 Hallucinatory proneness and age

Advancing age has been correlated with hallucinatory proneness ^55–57^, which may be associated with the development of age-related diseases that are paired with hallucinations ^55^. To test this in our data, we explored the relationship between age and our button press measures.

There was a positive correlation between the total proportional duration of Ganzfeld hallucinations and age (r_s(18)_=0.48, p=0.032, Supplementary Figure 3A) and the frequency of Ganzfeld hallucinations and age (r_s(18)_=0.51, p=0.022, Supplementary Figure 3B). No correlations (p<0.05) were found between the number, proportional duration or average duration of flicker hallucinations and age, or average duration of Ganzfeld hallucinations and age.

## 4 Discussion

Flicker and Ganzfeld are two visual stimulation methods that can generate visual hallucinations in people without pathological neurocognitive functioning and without the use of psychoactive pharmacological substances, and which differ in the degree of bottom-up visual input they involve. We compared the phenomenology of the hallucinations induced by these two methods to gain insight into the differences and commonalities of mechanisms giving rise to these hallucinations. Specifically, we predicted that flicker may elicit more simple hallucinations than Ganzfeld due to a relatively strong bottom-up signal (Hypothesis 1), and that there would be more simple hallucinations than complex hallucinations in flicker (Hypothesis 2). With regards to complex hallucinations, we considered two possible predictions: firstly, if both types of hallucinations rely on the spread of bottom-up activation sweeps, flicker might induce more complex hallucinations (Hypothesis 3a). Alternatively, if complex hallucinations rely strongly on top-down imagery-like processes, a Ganzfeld may elicit relatively more complex hallucinations due to the reduced sensory bottom-up input competing with this top-down drive (Hypothesis 3b).

As predicted, we found that flicker elicited more simple hallucinations than Ganzfeld (Hypothesis 1) corresponding to a stronger visual drive during flicker. We also found that both experimental conditions elicited substantially more simple hallucinations than complex hallucinations (Hypothesis 2). With regards to complex hallucinations, our results are largely consistent with Hypothesis 3a: there was a higher frequency of both simple and complex hallucinations in flicker compared to Ganzfeld, and the likelihood or duration of a complex hallucination relative to a simple one was not significantly higher during Ganzfeld than flicker. We also found a positive correlation between the total proportional time spent hallucinating in flicker and Ganzfeld hallucinations (Hypothesis 4), validated via the IEQ. This provides evidence for a common mechanism underlying flicker and Ganzfeld. Together, our data suggest that bottom-up visual signals could be a main driving factor for simple and complex hallucinations in both flicker and Ganzfeld.

There is a multitude of evidence to suggest that low-level aberrant noise, excitability, or direct activation of early visual cortices, which may be interpreted as signal, may be responsible for hallucinatory experiences. Patients with Charles Bonnet syndrome present with a build-up of neural activity in early visual cortex prior to the onset of a hallucination ^58^, and respond well to inhibitory stimulation of the visual cortices ^59^. Psychedelic drugs such as LSD and psilocybin may bring about some of their low-level geometric visual effects from direct activation of 5HTA receptors, which are populous in early visual cortex, causing excitation ^16^. Migraine sufferers, who may experience aura and hallucinations, are thought to have a hyperexcitable brain ^60^. It is possible that when the combined effects of steady internal activation and transient external stimulation exceed a certain threshold, pattern formation can occur^34^. For instance, sub-hallucinatory doses of mescaline, when combined with flicker, result in hallucinations above and beyond those normally seen in flicker ^34,61^. It is possible that fluctuations in neuronal excitability that are present in pathological and pharmacological states are responsible for hallucinatory proneness (to a lesser degree) during Ganzfeld. This internal neuronal excitability could act synergistically with the external input that flicker provides to result in the increased observed incidence of simple hallucinations in the flicker compared to the Ganzfeld.

Both experimental conditions elicited substantially more simple hallucinations than complex hallucinations (Hypothesis 2). This finding is consistent with complex hallucinations emerging from visual cortex excitation at higher stages of the visual system, i.e. a feedforward extension of simple hallucinations. There are unquestionably further factors beyond activation of early visual cortices which may predispose an individual to experience complex hallucinations (such as personality traits like absorption ^18,46^, strength of mental imagery ^14,42^ and positive schizotypy ^48^). Each of these factors may act independently of visual stimulation technique and the tendency to experience simple hallucinations, given the higher frequency of simple hallucinations in both tasks, and would likely be less effective at altering the tendency to experience simple hallucinations given their greater dependence on earlier visual areas.

With regards to the mechanisms underlying simple vs. complex hallucinations, our results are broadly consistent with Hypothesis 3a: that the mechanisms underlying simple and complex hallucinations are the same, but are driven more strongly during flicker. Namely, there was a higher frequency of both simple and complex hallucinations in flicker compared to Ganzfeld, and the likelihood of a complex hallucination relative to a simple one was not significantly higher during Ganzfeld than flicker. It should be noted that our sample was relatively small (n=20), and we used subjective, researcher-defined criteria to determine the complexity of hallucinations (Supplementary Materials, Table 1), so it is possible that variations in hallucination complexity in line with a different balance in bottom-up and top-down processes across the Ganzfeld and flicker techniques will be found with larger samples or different criteria. We also note that we found an interaction effect between condition and complexity in our analyses of the IEQ questionnaire (though not the ASC), showing that the difference between Simple and Complex Imagery scores was greater for the flicker than the Ganzfeld condition. This provides some evidence for Hypothesis 3b - that reduced visual input during sensory deprivation may allow for more complex imagery-like hallucination content. However, the lack of such an interaction on all other measures, alongside the non-significance of a supplementary Chi-squared test, means that any such shifts in the balance between simple and complex hallucinations across these two methods are likely to be quite subtle.

Button presses provide a direct measure of hallucinations but might be influenced by factors not directly related to phenomenology, such as the criterion for when to press. Furthermore, the classification of hallucinations indicated by button presses and prompts as simple or complex relied on subjective judgments by the researchers. Hypothetically, the criteria used to define complexity may have differed between experimenters and participants. A crucial aspect of our study was therefore that the results from these button press and prompting responses were largely replicated by independent retrospective questionnaire measures. This, alongside the absence of correlations between button press measures and participants’ perception of button press interference, suggests that our approach is appropriate for quantifying hallucinatory experience without significantly impacting participants’ overall experience. Ultimately, self-report measures are, by nature, subjective, difficult to verify and can be influenced by various factors, and this is a major challenge in the study of hallucinations. In future, the use of additional objective measures, such as neuroimaging techniques, could provide more objective and detailed information about the neural correlates of hallucinations and their phenomenology induced by different stimulation methods.

We show descriptively that the time course of the hallucinations varied across conditions, with complex hallucinations demonstrating a delayed onset compared to simple hallucinations in both the flicker and the Ganzfeld. Interestingly, during flicker, complex hallucinations peaked shortly after simple hallucinations, whereas in the Ganzfeld, complex hallucinations took substantially longer than simple hallucinations to peak. One possible explanation within the framework of our hypotheses could be that the stronger visual signals in the flicker cause hallucination-inducing activation to spread more rapidly through the hierarchy than the weaker noise fluctuations in the Ganzfeld. Interestingly, we observed a somewhat bimodal distribution of simple hallucinations among participants in both the flicker and Ganzfeld conditions. This could be due to inhomogeneous and potentially cyclical activation dynamics over time, which could be an interesting hypothesis to explore in future studies of an extended duration.

A deeper explorative analysis of hallucination content as indicated by words used to describe the experience and associated drawings, revealed a greater presence of form constants in flicker compared to Ganzfeld. During flicker, verbal descriptions and drawings often contain references to potential Klüver-like forms such as ‘spirals’, ‘tunnels’, ‘patterns’ and ‘spinning’. Surprisingly, these references were absent in the Ganzfeld condition, raising the possibility that the percepts seen during Ganzfeld may bear a closer resemblance to phosphenes. It is also worth considering the relative uncertainty associated with the words used to describe the Ganzfeld condition (for instance shadows, faded, disappeared, slightly, sort). This may suggest that some percepts in Ganzfeld lack the clarity, spatial resolution, and vividness observed in the largely bottom-up driven flicker hallucinations ^24,26,62,63^.

An unexpected finding was the increased duration of Ganzfeld hallucinations compared to flicker hallucinations. This may be related to the temporal modulatory input that flicker provides, which may contribute to the instability of the hallucinatory state, as each temporal modulation acts as not only a stimulus to pattern formation, but also an agitation of previously elicited patterns ^34^. The absence of these disruptions in the Ganzfeld would likely promote longer hallucinations. Of relevance is that both flicker and Ganzfeld were associated with words of dynamic nature, evidenced in frequency analyses and word clouds, which highlight key terms such as spinning, movement, and changing – perhaps even to a higher degree during flicker (Table 1).

A positive correlation was found between age and the number and proportional duration of Ganzfeld hallucinations. Advancing age has been correlated with hallucinatory proneness ^55–57^. One explanation for these findings is the excitability of visual cortex. As previously mentioned, multiple sources of evidence suggest that excitability of early visual cortices may be responsible for or precede hallucinatory experiences ^3,45,58–60^. While more research in larger samples is needed to test this hypothesis, it is plausible that age-related changes in the excitability of the visual system could occur, contributing to the generation and propagation of hallucinatory experiences. These changes could particularly impact hallucinatory experience in the Ganzfeld, as any changes in excitability resulting in hallucinatory experience may be more dependent on individual variability in excitability without strong bottom-up visual drive. However, studies on cortical excitability of the motor cortex in the ageing brain have generally reported reduced excitability^64^, so more research on this process in the visual domain is needed. It is also possible that increased hallucination proneness with age relies on a different process. Predictive coding accounts, for example, have suggested age-related upweighting of predictions about the world ^25,65–67^.

Ultimately, this study lays the groundwork for further investigations into the characteristics and mechanisms of hallucinations induced by different stimulation techniques. Building upon the findings of this study, future research could address several important avenues of inquiry. It would be valuable to further explore the relationship between baseline behavioural measures (such as mental imagery^14,42^ and positive schizotypy^68^) and the propensity to experience complex hallucinations across various stimulation methods. Understanding how individual differences influence hallucination generation could contribute to personalized approaches in hallucination research and potentially inform clinical interventions for individuals experiencing pathological hallucinations. Moreover, investigating the neurobiological mechanisms underlying hallucinations induced by flicker and Ganzfeld stimulation could provide a deeper understanding of the neural processes involved in perceptual abnormalities. Neuroimaging and brain stimulation could be used to explore whether more excitable individuals are more prone to visual hallucinations in flicker and Ganzfeld, either by looking at cortical excitability at baseline ^69^ or during the specific time course of discrete hallucinatory periods ^58^.

In conclusion, this study provides insights into the phenomenology of hallucinatory experiences and their relationship with different stimulation techniques. We find evidence to suggest that the mechanisms underlying simple and complex hallucinations in flicker and Ganzfeld are largely driven by a bottom-up mechanism, which is amplified in flicker compared to Ganzfeld due to heightened bottom-up input.

## Data availability statement

Data is available on request.

## Authorship contribution statement

OS: Conceptualisation, Methodology, Data collection, Analysis, Visualisation, Writing – Original draft preparation. ML: Analysis, Writing – Reviewing and editing. JAG: Supervision, Writing – Reviewing and editing. JIS: Conceptualisation, Methodology, Supervision, Writing – Reviewing. TMD: Conceptualisation, Methodology, Supervision, Writing – Reviewing and editing. All authors reviewed and approved the final manuscript.

