## Supplementary Materials for "Flicker and Ganzfeld induced visual hallucinations differ in frequency and content"

*Detailed participant instructions*

Prior to the experiment, participants were given brief examples of the types of visual experiences they might experience they were informed that they could see any combination of a) simple or abstract hallucinations such as colours, movement, patterns such as kaleidoscopes, or shapes or b) complex or figurative hallucinations, such as objects, animals, faces or scenes. They were also told that they may not see anything at all. They were informed that these visual experiences were of a 'pseudo-hallucinatory' nature, and that they would be aware that they were seeing was not real. They were encouraged to be patient while waiting for any experiences to arise, especially with the Ganzfeld condition. Participants were asked not to engage in any active daydreaming or mental imagery, and to passively watch any observations, making sure to pay attention to them and allow them to evolve if necessary. To this end, participants were asked not to verbalise their hallucinations until the experience faded away, however, were not interrupted if they began verbalising their experience while the hallucination was ongoing. Participants were instructed to keep their eyes open for both conditions.

Monitoring of, and communication with participants during the flicker and Ganzfeld hallucination phases of the study took place via video and audio call from a nearby room. Communication was kept to a minimum to avoid interruption of the hallucinatory state. In some cases, the experimenter was present within the room, not in eyesight of the participant. This did not have an effect any measures collected. After each hallucination condition, retrospective measures of experience (drawings, interviews, and questionnaires) were completed in a separate room.

*Hallucination classification*

**Supplementary Table 1:** Examples of prompts, drawings, and hallucination classification

| Prompt | Drawing | Hallucination classification |
|---|---|---|
| Spinning kaleidoscope at the bottom, like a tornado | 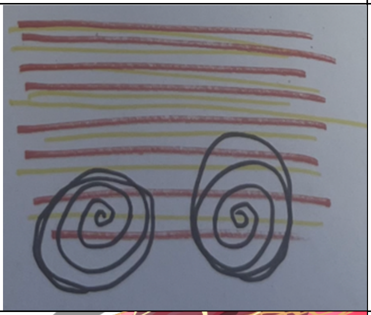 | Simple |
| Penguin merged with square shapes | 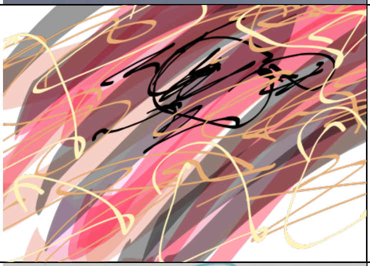 | Complex |
| Big wave, dark blue behind it | 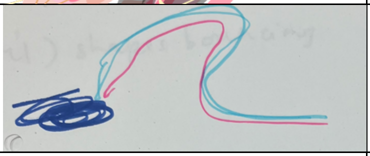 | Simple |
| Indistinct dog | 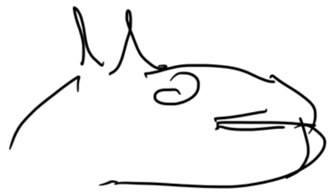 | Complex |
| Random shapes | 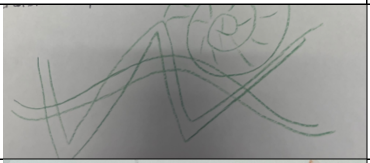 | Simple |
| Ring around a planet round a shape diagonally spinning, galaxy | 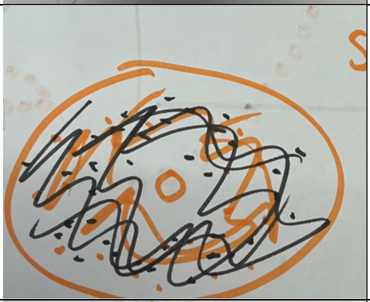 | Complex |
| Tunnel goes away like Mario-Kart, rainbow road | 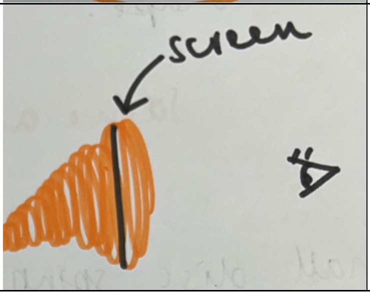  | Simple |

| | | |
|---|---|---|
| Very faint skull | 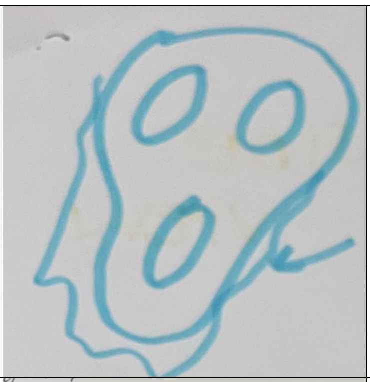 | Complex |
| Whispy octopus legs | 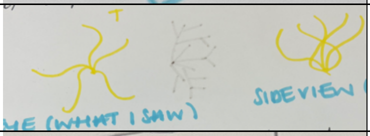 | Simple |
| White door | 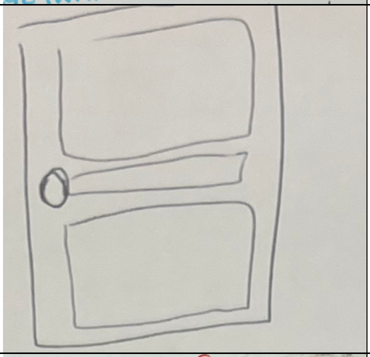 | Complex |
| Introduction to Dr Who | 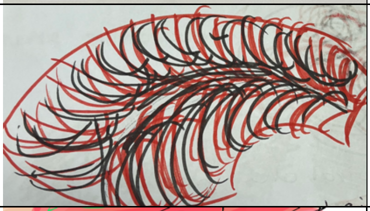 | Simple |
| Tortoises | 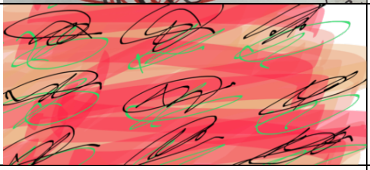 | Complex |
| Snowflake | 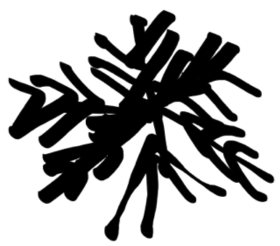 | Simple |
| Microbe spots moving into a centre point | 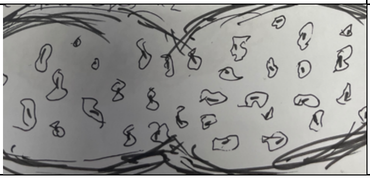 | Complex |

| | | |
|---|---|---|
| Ball rolling in the middle of the screen | 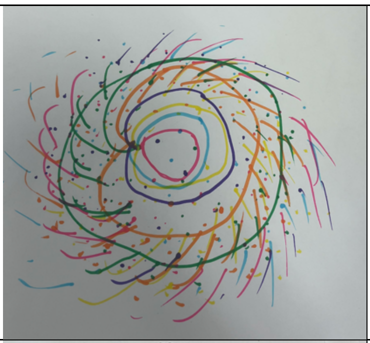 | Simple |
| Symmetrical face turning left to right | 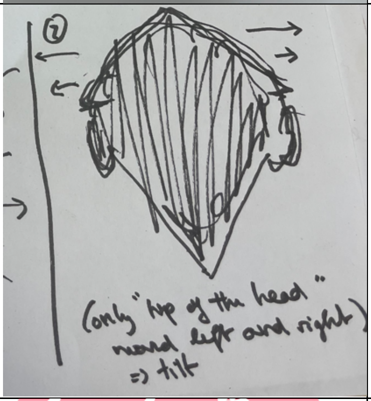 | Complex |
| Giant ball on the centre of the screen keeps vibrating | 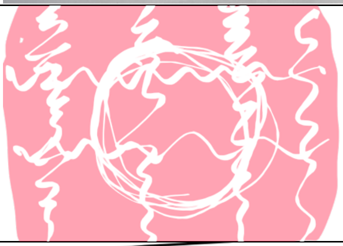 | Simple |
| Mirror image of a butterfly | 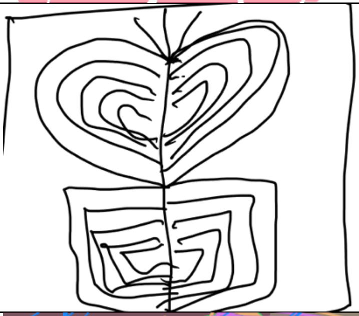 | Complex |
| Green electric currents in middle expanding and merging with orange, elements of blue, purple, blue dots | 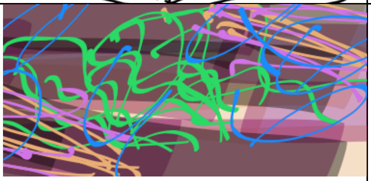 | Simple |
| One single caveman face | 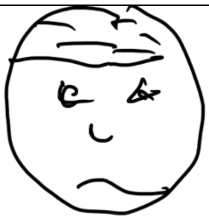 | Complex |

*Questionnaires*

The questions comprising the ASC-R can be analysed by dissecting the components into five primary dimensions (5D-ASC - Auditory Alteration, Dread of Ego Dissolution; Oceanic Boundlessness; Reduction of Vigilance; Visionary Restructuralisation) or 11 primary dimensions (11D-ASC - Anxiety; Audio-Visual Synaesthesia; Blissful State; Changed Meaning of Percepts; Complex Imagery; Disembodiment; Elementary Imagery; Experience of Unity; Impaired Control and Cognition; Insightfulness; Spiritual Experience). Specifically, we used questions from the 11D components Elementary Imagery (items 14 and 22). We did not use item 33 from the Elementary Imagery component (I saw lights or flashes of light in total darkness or with closed eyes), due to this being reflected in the nature of the stimuli. We also used questions from the Complex Imagery component (items 39, 72 and 82). Many of these questions were followed by the statement 'in complete darkness or with closed eyes'; this statement was removed for the purpose of this study as there was always some visual input, and our participants were asked to keep their eyes open. We also utilised one question from the Positive Derealisation subscale of the Oceanic Boundlessness component (5D, item 1) and one question from the visionary reconstruction component (5D, item 7). In addition, a catch question was used (a repetition of item 7) to ensure participants were paying attention to the question.

**Supplementary Table 2**: Items from ASC-R utilised in study

| Dimension | Item | Question |
|---|---|---|
| Elementary Imagery (11D) | 14 | I saw regular patterns [in complete darkness or with closed eyes] |
| | 22 | I saw colours before me [in total darkness or with closed eyes] |
| Complex Imagery (11D) | 39 | I saw scenes rolling by [in total darkness or with my eyes closed] |
| | 72 | I could see pictures from my past or fantasy extremely clearly |
| | 82 | My imagination was extremely vivid |

| | | |
|---|---|---|
| Oceanic Boundlessness (5D) | 1 | I felt like I was in a fantastic other world |
| Visionary Reconstruction (5D) | 7 | I saw things that I knew were not real* |

*Question repeated as a catch question. Participants were scored on a percentage difference between a repetition of the catch question (I saw things that I knew were not real) for both hallucinatory conditions. A percentage difference of greater than 30% between the catch and the original question in both conditions was used as criterion for exclusion for the participant. No participants met this criterion.

**Supplementary Table 3:** Items from IEQ utilised in current study

| Dimension | Item | Question |
|---|---|---|
| Complexity | 1 | I saw bursts of light or splashes of colour. |
| | 2 | I saw abstract geometrical designs and patterns. |
| | 3 | I saw rapidly transforming objects/ figures. |
| | 4 | I saw repetitive, moving objects/ figures embedded in geometrical patterns. |
| | 5 | I saw stable, well-defined objects/ figures. |
| | 6 | I saw snapshots or glimpses of full scenes |
| | 7 | I saw full-fledged scenes without being a part of them, similar to watching a movie |
| | 8 | I was fully immersed within what looked and felt like another authentic realm |
| | 9 | I was surrounded by a supreme white light |
| Progressive | 17 | My vision progressed over time from simple (busts/splashes/geometries) to complex (well-defined objects/figures) images. |
| | 18 | My vision progressed over time from isolated elements to full, immersive scenes. |

A

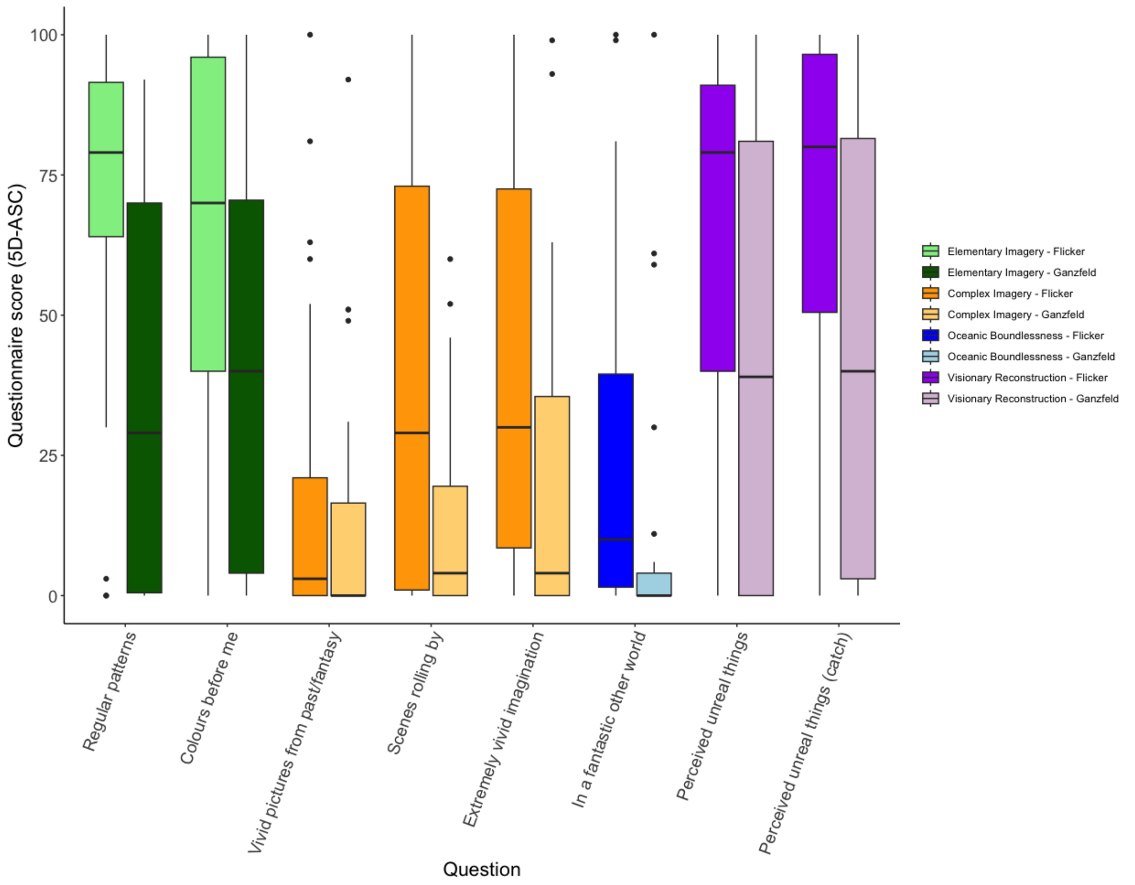

Questionnaire score (5D-ASC)
100
75
50
25
0
Regular patterns
Colours before me
Vivid pictures from past/fantasy
Scenes rolling by
Extremely vivid imagination
In a fantastic other world
Perceived unreal things
Perceived unreal things (catch)
Question
Elementary Imagery - Flicker
Elementary Imagery - Ganzfeld
Complex Imagery - Flicker
Complex Imagery - Ganzfeld
Oceanic Boundlessness - Flicker
Oceanic Boundlessness - Ganzfeld
Visionary Reconstruction - Flicker
Visionary Reconstruction - Ganzfeld


B

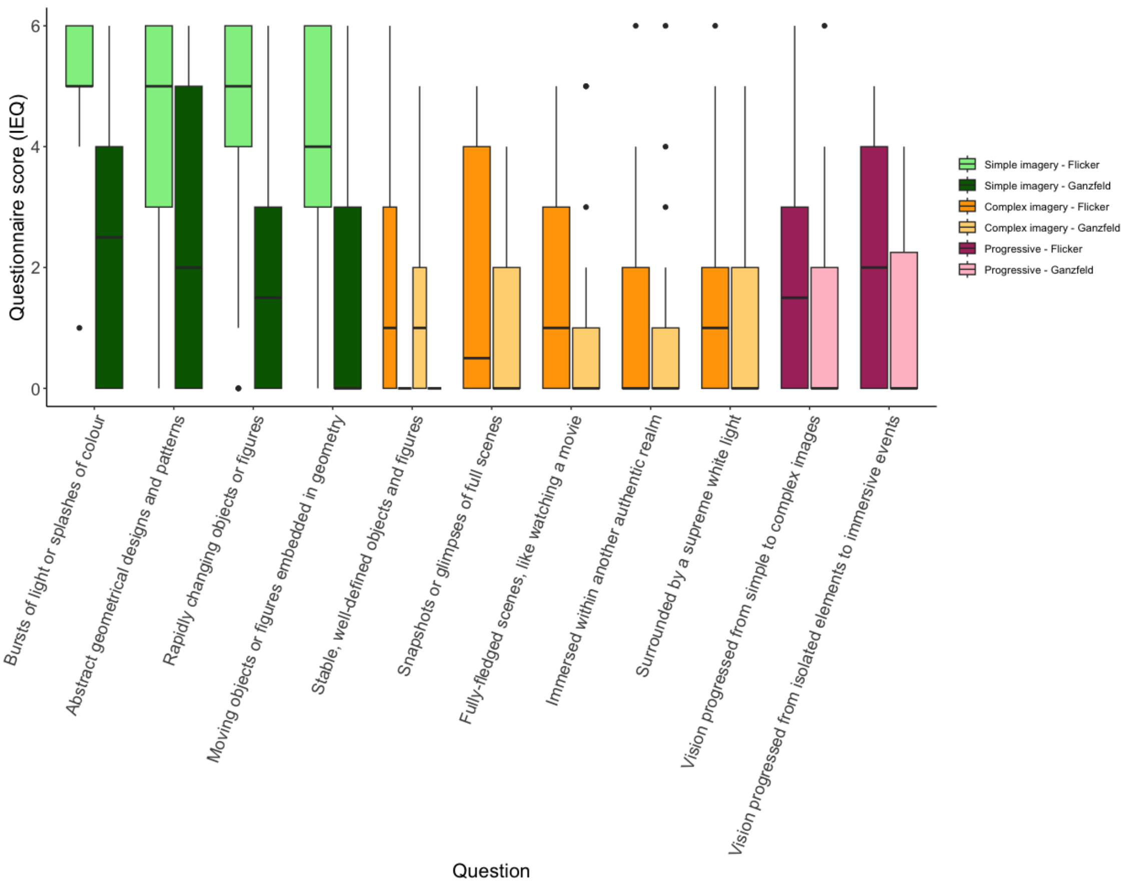

Questionnaire score (IEQ)
6
4
2
0
Bursts of light or splashes of colour
Abstract geometrical designs and patterns
Rapidly changing objects or figures
Moving objects or figures embedded in geometry
Stable, well-defined objects and figures
Snapshots or glimpses of full scenes
Fully-fledged scenes, like watching a movie
Immersed within another authentic realm
Surrounded by a supreme white light
Vision progressed from simple to complex images
Vision progressed from isolated elements to immersive events
Question
Simple imagery - Flicker
Simple imagery - Ganzfeld
Complex imagery - Flicker
Complex imagery - Ganzfeld
Progressive - Flicker
Progressive - Ganzfeld

**Figure 1**: (A) ASC questionnaire results separated out by question. Questions are split into separate dimensions (Green – Elementary Imagery; Orange – Complex Imagery; Blue – Oceanic Boundlessness; Purple – Visionary Reconstruction), with the darker shade of each question indicative of the flicker (and the lighter, Ganzfeld), (B) IEQ questionnaire results separated out by question. Questions are split into separate dimensions (Green – Simple Imagery; Orange – Complex Imagery; Pink – Progressive Imagery), with the darker shade of each question indicative of the flicker (and the lighter, Ganzfeld), Questions are abbreviated; please see Supplementary Tables 2 and 3 for unabbreviated questions. Both conditions are *N*=20, for two conditions. ASC – Altered States of Consciousness (rating scale); IEQ – Imagery Experience Questionnaire

.

*Button press interference and sleepiness*

Paired Wilcoxon rank signed tests were carried out to see how participants sleepiness and their perception of how much the button press interfered with their experience varied between the Ganzfeld and the flicker condition. Two participants were removed from the sleepiness analyses as their data was not recorded for one condition, leaving n=36 (as their data for their other condition was also excluded). Five further participants were excluded from the button press analyses as they had no hallucinations in one condition, therefore they could not provide an opinion on how much they thought the button presses interfered with their experience, resulting in n=26 (as their data for their other condition was also excluded).

Paired Wilcoxon signed rank tests suggested a difference between participants sleepiness in the flicker condition compared to the Ganzfeld condition (V=0, $p<0.001$). There was no difference between how much participants perceived the interference of the button press in either condition (V=33, p=1).

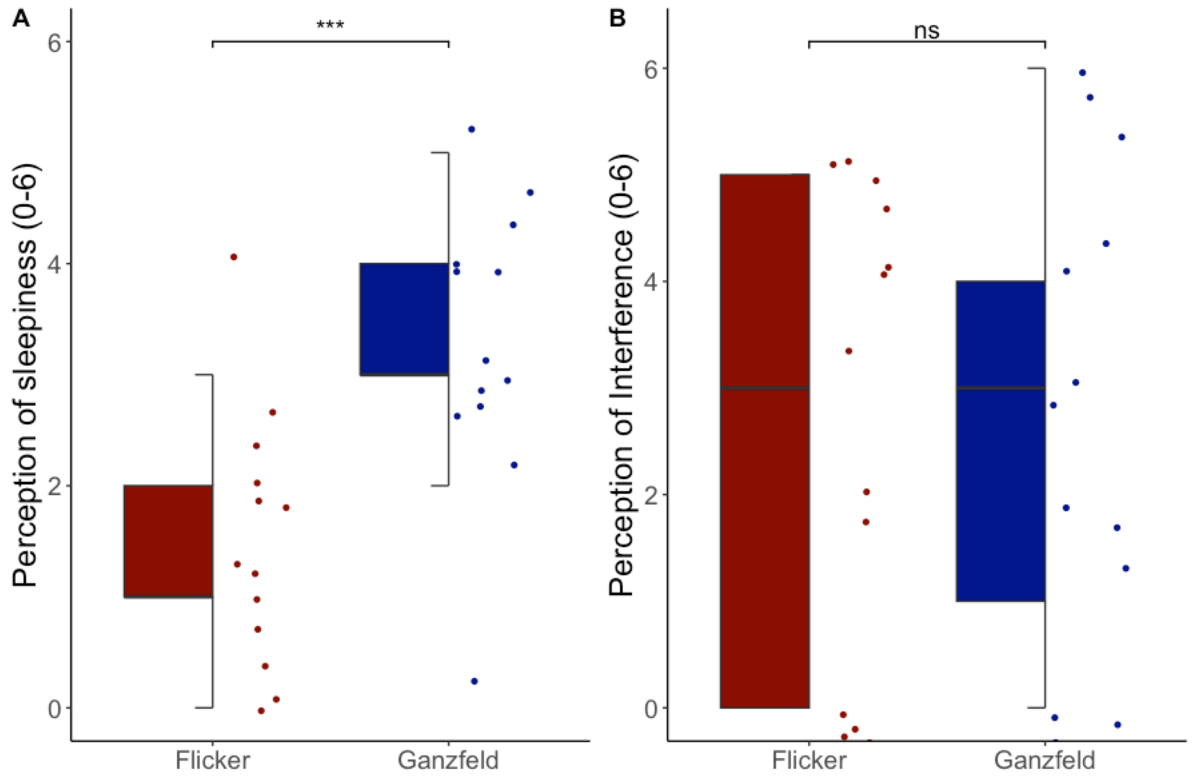


**Figure 2: (A)** Relationship between experimental condition (x-axis: red – flicker; blue – Ganzfeld) and participants perception of sleepiness (y-axis) (B) Relationship between experimental condition (x-axis; red – flicker; blue – Ganzfeld) and participants perception of how much the button press interfered with their hallucinatory experience (y-axis). Statistics shown are Paired Wilcoxon rank signed tests (*N*=20)

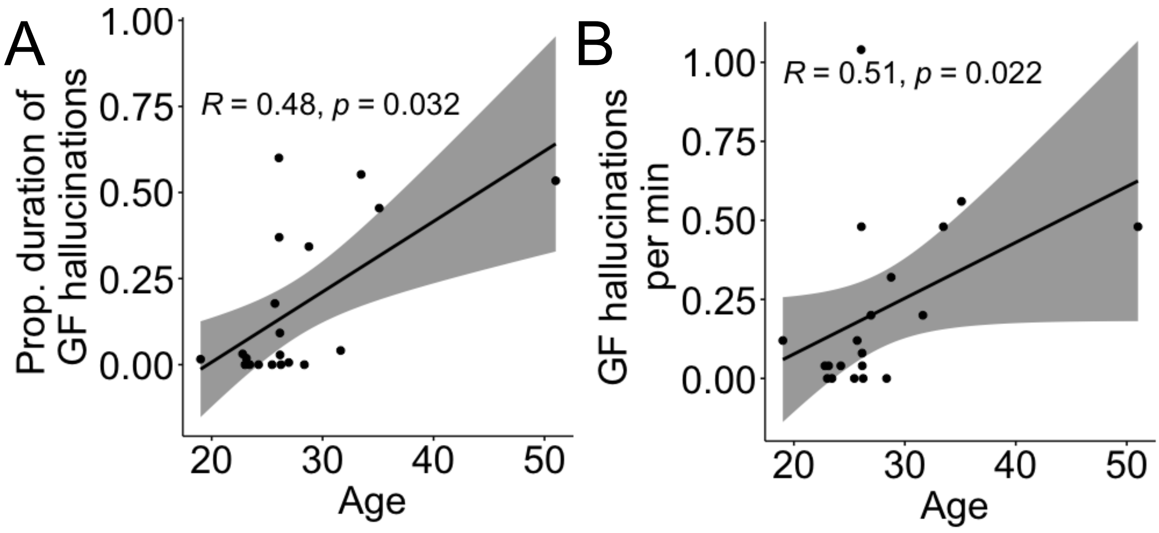


**Figure 3:** (A) Scatter plots showing the relationship between the total proportional time spent hallucinating during the Ganzfeld (y-axis) and age (x-axis) (B) Scatter plots showing the relationship between Ganzfeld hallucinations per minute (y-axis) and age (x-axis). Both plots show associated trend line (black) and Spearman's rank correlation testing (95% CI; grey shading) for *N*=20. GF – Ganzfeld; min – minute; prop – proportional.
